# Evidence for microbially-mediated tradeoffs between growth and defense throughout coral evolution

**DOI:** 10.1101/2023.04.26.538152

**Authors:** Hannah E. Epstein, Tanya Brown, Ayomikun O. Akinrinade, Ryan McMinds, F. Joseph Pollock, Dylan Sonett, Styles Smith, David G. Bourne, Carolina S. Carpenter, Rob Knight, Bette L. Willis, Mónica Medina, Joleah B. Lamb, Rebecca Vega Thurber, Jesse R. Zaneveld

## Abstract

Evolutionary tradeoffs between life-history strategies are central to animal evolution. However, because microbes can influence aspects of host physiology, behavior, and resistance to stress or disease, changes in animal-microbial symbioses have the potential to mediate life-history tradeoffs. Scleractinian corals provide a highly biodiverse and data-rich host system to test this idea, made more relevant by increases in coral disease outbreaks as a result of anthropogenic changes to climate and reef ecosystems. Identifying factors that determine coral disease susceptibility has therefore become a focus for reef conservation efforts. Using a comparative approach, we tested if coral microbiomes correlate with disease susceptibility across 425 million years of coral evolution by combining a cross-species coral microbiome survey (the “Global Coral Microbiome Project”) with long-term disease prevalence data at multiple sites. Interpreting these data in their phylogenetic context, we show that microbial dominance and composition predict disease susceptibility. We trace this dominance-disease association to a single putatively beneficial bacterial symbiont, *Endozoicomonas*, whose relative abundance in coral tissue explained 30% of variation in disease susceptibility and 60% of variation in microbiome dominance across 40 coral genera. Conversely, *Endozoicomonas* abundances in coral tissue strongly correlated with high growth rates. These results demonstrate that the evolution of microbial symbiosis in corals correlates with both disease prevalence and growth rate. Exploration of the mechanistic basis for these findings will be important for our understanding of how microbial symbiosis influences animal life-history tradeoffs, and in efforts to use microbes to increase coral growth or disease resistance *in-situ*.

**Significance Statement:** The evolution of tropical corals, like that of many organisms, involves tradeoffs in life-history strategy. We sought to test whether microbes influence coral life-history traits. Comparative data from a census of modern coral microbes, combined with long term disease surveys in three regions, provide evidence for a correlation between microbiome structure, growth rate, and disease susceptibility during coral evolution. These trends were driven primarily by changes in the relative abundance of *Endozoicomonas* in coral tissue microbiomes, suggesting the novel hypothesis that *Endozoicomonas* may allow corals to grow more quickly at the cost of greater vulnerability to disease. Thus, symbiosis with microbes may be an important aspect of animal life-history strategy.

## Introduction

Tradeoffs in life-history strategy are key features in animal evolution (1, 2). These tradeoffs often involve differential investments in life-history traits such as growth rate (3); reproductive maturation, timing, and fecundity (4); or resistance to stress (5), predation (6) or disease (7). The fitness costs and benefits of these investments are often context-dependent. Thus, shifts in ecological or environmental conditions can favor some life-history strategies over others (5), sculpting trait evolution within animal lineages and reshaping ecological communities. Global climate change is shifting the patterns and prevalence of disease in many animal taxa, while increasing the virulence of some pathogens (8, 9). Identifying evolutionary tradeoffs and resulting trait correlations associated with disease susceptibility (10) can therefore help predict how species survival will shift with climate change.

Although much research on evolutionary tradeoffs focuses on the traits of animals themselves, it is also well documented that the physiology (11), fitness and even behavior (12) of many animals are influenced by their microbiomes. Ecological microbiome surveys and laboratory experiments using germ-free animals have linked animal microbiomes, and specific symbionts within them, to multiple key life-history traits, including growth (13), development rate (13), fecundity (13), stress resistance (11, 14), and disease susceptibility (14). It therefore seems likely that microbial symbiosis is an important aspect of animal life-history tradeoffs.

If microbes do influence life-history traits (or vice versa), microbiome structure and membership may correlate with those traits over long periods of animal evolution. However, testing the potential relevance of microbial symbiosis for life-history strategy over evolutionary time periods is challenging. These tests must use phylogenetic comparative methods that account for trait correlations induced by the shared history of traits over evolution. They further require large cross-species datasets on both animal traits and microbiome structure. Scleractinian corals meet these data requirements and are therefore an animal lineage that present a unique opportunity to answer the question of whether microbes and life-history strategies are associated.

The reef-building corals that have evolved over 425 million years represent a diverse group of animals, including an estimated >1600 species (15), with an extensive fossil record, and a well-known variety in both life-history strategy (2) and microbial symbiosis (16–18). These animals also have special ecological and societal importance, as corals are foundational to reef ecosystems that support some of the most biodiverse assemblages on the planet. These ecosystems in turn support the livelihoods of the many coastal communities that rely on them for food, coastal protection, and recreation (19). Yet the ancient diversity of coral reefs is currently threatened by global climate change, which is driving both dramatic mass bleaching events and increased prevalence and severity of disease outbreaks (8). Due to the threats to coral reef ecosystems, and the potential harm that their collapse could inflict on millions of people in coastal communities worldwide, corals and their microbiomes have been intensively researched.

Association with specific dinoflagellate symbionts (e.g., *Duruisdinium* vs. *Cladocopium*) has been reported to have complex and species-specific influences on coral traits such as thermal tolerance and growth rate ((20) but see (21)). Evolutionary studies have demonstrated that vertical vs. horizontal transmission of these dinoflagellate symbionts is tied to important host traits, such as reproduction by brooding vs. spawning (22). However, potential influences of symbiosis with bacteria and archaea on coral life-history traits are less well understood.

Coral microbiome research has demonstrated that in the present-day communities of coral-associated bacteria and archaea (hereafter ‘coral microbiomes’) are influenced by host traits; local environmental factors, such as temperature, depth, nutrient availability, and turbidity; anatomy (16); and ecological context, such as predation, exploitation by farming fish (23), or competition with turf algae. Specific microbes have been shown to protect corals from pathogens through antimicrobial production (24), predation (25), jamming of quorum-sensing systems (26), and passive competition for space and resources. Differences in microbiome structure or dynamics are also often found between related species that show different patterns of disease susceptibility (27). These examples provide support for connections between coral life-history, microbiome structure and disease susceptibility in the present day, although they do not directly allow for statistical testing of evolutionary hypotheses.

Clarifying whether microbiome structure and coral life-history traits correlate over coral evolution globally would provide vital context for interpreting studies of extant coral symbiosis and disease at local or regional scales. Several lines of research have created a strong foundation on which such comprehensive comparative evolutionary analyses can be built. Coral disease patterns have been intensively researched, and an increasing number of datasets are now openly available. Well-curated global databases of coral physiological traits (28), with contributions from numerous research groups, have been established and mapped to coral life-history strategies (2). Finally, several large cross-species studies of corals and their microbiomes have been launched. These advances provide an opportunity to compare host trait data and microbiome structure from across the coral tree of life.

Here, we test whether microbiome structure correlates with coral disease susceptibility, growth rate, or overall life-history strategy. To address this question quantitatively, we first characterized the microbiome composition from visibly healthy samples of 40 coral genera using 16S rRNA gene sequencing results from the Global Coral Microbiome Project (16)(Supplementary Data Table 1a), and subsequently combined these data with genus-level long-term disease prevalence data from several tropical regions around the globe (the Caribbean (Florida Reef Resilience Project data (FRRP, https://frrp.org/)), central Pacific (Hawai’i Coral Disease Database (HICORDIS) (29)), and eastern Australia (this study); Supplementary Data Table 1b), and coral life-history traits from the Coral Trait Database (28) (Figure 1). With the resulting microbiome structure, disease prevalence, and coral growth data across a global distribution of coral genera (Supplementary Data Table 1c), we compared these traits using methods that account for phylogenetic correlations using a time-calibrated multi-gene reference tree of corals (30).

**Figure 1.**
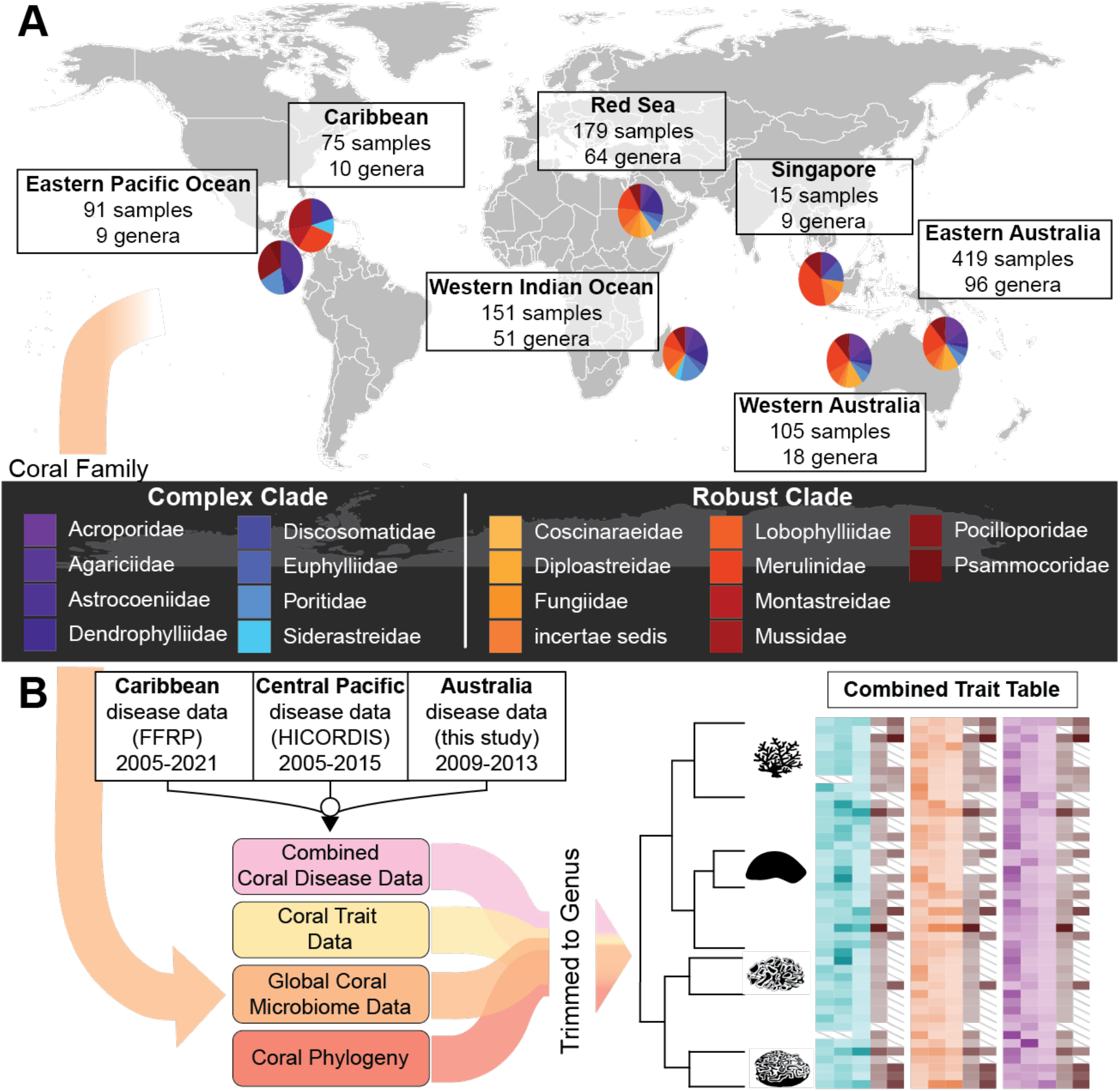
Conceptual overview of data sources integrated for the project. **A**. Map of sampling locations for coral microbiomes analyzed in the manuscript. Pie charts show the proportion of coral samples from families in the Complex clade (cool colors) and Robust clade (warm colors). Samples were collected from coral mucus, tissue, and endolithic skeleton (see Methods). **B**. Schematic representation of data integration for the project. Coral microbiome data (as shown in A) were combined with long-term disease prevalence data from 3 projects (the Florida Reef Resilience Program (FFRP), the Hawai’i Coral Disease Database (HICORDIS), and data from Australia (this study)), as well as coral trait data from the Coral Trait Database, and a molecular phylogeny of corals (see Methods). In order to integrate data from these disparate sources, all annotations were pooled at the genus level. The end product is a trait table of microbiome, taxonomic, physiological, and disease data across diverse coral genera.

Across coral evolution, we found that microbiome structure in healthy corals is correlated with both disease-susceptibility and growth rate. We further identified these correlations as being primarily driven by a single key bacterial genus, *Endozoicomonas*, a common coral symbiont that often forms aggregates within coral tissue (31) and is hypothesized to be a metabolic mutualist. These results provide an important example of long-term correlations between microbiome structure and host traits (disease susceptibility and growth rate), supporting the notion that microbial symbiosis can have important roles in mediating animal life-history tradeoffs.

## Results and Discussion

### Coral microbiomes are dominated by a small number of bacterial taxa

The microbiome of corals is often dominated by a few highly-abundant taxa that demonstrate species-specificity (17, 18), though why these highly-abundant microbial taxa differ across coral diversity is unknown. To test this, we first identified a restricted set of dominant bacterial or archaeal taxa in visibly healthy corals retrieved from mucus, tissue, and skeleton samples of 40 coral genera. (‘Dominant taxa’ were defined as those that are most abundant on average within all samples from a given portion of coral anatomy in a given coral genus.) Thirty-eight of the coral genera were dominated by the bacterial classes α- or γ-proteobacteria, which are are known to include common coral associates (17), with further detailed taxonomic resolution revealing that the number of dominant bacterial and archaeal genera across compartments also remained limited (Figure 2A; Supplementary Data Table 2). For example, only 17 genera of bacteria or archaea accounted for the dominant microbes in the tissue microbiomes of all 40 coral genera (this number excludes 4 unclassified ‘genera’ that could not be classified to at least the order level). Mucus and skeleton showed similar trends, with only 16 and 25 dominant genera, plus 2 or 4 unclassified genera, respectively. Across coral genera, *Pseudomonas* was most commonly dominant in mucus (31.4% of coral genera), while *Endozoicomonas* was most commonly dominant in tissue (18%) and *Candidatus* Amoebophilus (13.5%) was most commonly dominant in skeleton microbiomes. Currently the influences of microbiome structure and dominance of particular microbial taxa on coral physiology are not yet well understood.

**Figure 2.**
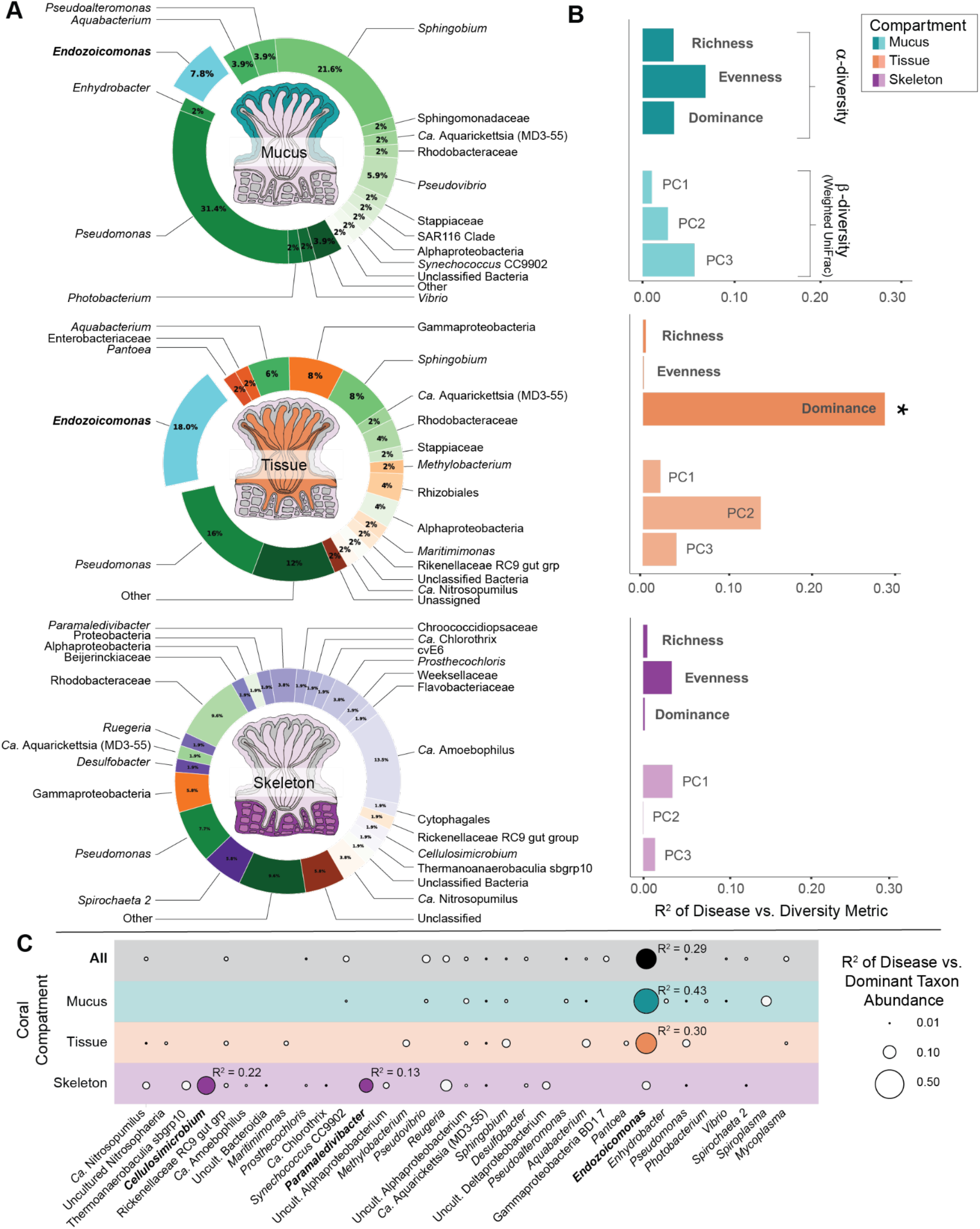
Dominant microbes in the coral microbiome. **A)** Dominant bacterial or archaeal genera in coral mucus (cyan), tissue (orange), or skeleton (purple) microbiomes. Pie wedges represent the fraction of coral host genera in which the labeled bacterium is more abundant than all other bacterial or archaeal taxa. Cyan shades represent microbes dominant in mucus, oranges represent microbes dominant in tissue (but not mucus), purple shades represent microbes dominant in skeleton (but not mucus or tissue). *Endozoicomonas*, which is of special significance later in the paper, is highlighted in aqua. **B)** Bar charts showing correlations between microbiome alpha and beta diversity metrics and disease, represented by the R^2^ for PGLS correlations. Alpha diversity metrics include richness, evenness (Gini index), and dominance (Simpson’s index), and weighted UniFrac beta diversity metrics including the three principal component axes (PC1, PC2, PC3) that represent measures of community structure. Significant relationships (p < 0.05, Supplementary Data Table 4) are marked by an asterisk (*). **C)** Bubble plot showing correlations between dominant microbial taxa and coral disease prevalence. The size of each bubble represents the R^2^ for PGLS correlations between disease susceptibility and microbial relative abundance for each listed taxon in either all samples (top row), mucus samples (cyan row), tissue samples (orange row), or skeleton samples (purple row). Colored points were significant (p < 0.05; Supplementary Data Table 6a). Points that were not significant or had too little data (n < 10) for reliable testing are marked in white. Taxa whose abundance is significantly correlated with disease are marked in bold on the x-axis.

### Microbiome richness and evenness do not predict disease susceptibility

To identify how bacterial communities are structured among globally distributed coral taxa, we characterized alpha diversity within the mucus, tissue, and skeleton compartments for each coral genus using several metrics. These included observed richness; the Gini index, which measures evenness. We visualized the evolution of each of these measures of microbiome alpha diversity using ancestral state reconstruction (Supplementary Figures 1a and 1b), then compared them against disease susceptibility using Phylogenetic Generalized Least Squares (PGLS) analysis. While we hypothesized that coral microbiomes high in overall biodiversity might show reduced disease susceptibility — analogous to the ability of more biodiverse ecosystems to resist invasive species (32) — neither microbiome richness nor evenness were significantly correlated with host disease susceptibility in phylogenetic generalized least squares analysis (PGLS richness vs. disease susceptibility: R^2^ = 0.004, *p* = 0.674, FDR q = 1; PGLS evenness vs. disease susceptibility R^2^ = 0.028, *p* = 0.274, FDR q = 1; Supplementary Data Table 3a). Some specific cases of coral genera with low microbiome richness and high disease susceptibility were identified (i.e., *Pocillopora, Acropora*, and *Montipora*; Supplementary Figure 1a & b) but there was no overall trend across all genera surveyed (Figure 2B). Thus, microbiome richness or evenness alone does not predict coral disease susceptibility.

### Microbiome dominance correlates with coral disease susceptibility

Given that neither microbiome richness nor evenness significantly predicted disease susceptibility, and that cross-species differences in a limited number of dominant microbes were very notable in the data, we hypothesized that corals with highly abundant bacterial taxa might display more disease vulnerability. To quantify this, ecological dominance among identified ASVs was calculated using Simpson’s Index, which estimates the probability that two species drawn from a population belong to the same group, and thereby incorporates aspects of both richness and evenness simultaneously. We correlated Simpson’s Index against coral disease prevalence for either all coral samples, or those in mucus, tissue, or skeleton considered individually. In coral tissue, microbiome dominance significantly correlated with disease, explaining roughly 27% of overall variation in disease susceptibility across coral species (PGLS: R^2^ = 0.27, p = 0.0006, FDR q = 0.025; Supplementary Data Table 3a; Supplementary Figure 1c). No other combination of alpha diversity measure and compartment correlated with disease after accounting for multiple comparisons (Figure 2B). Thus, microbiome dominance as measured by Simpson’s Index was a far stronger predictor of coral disease susceptibility than α-diversity measures that considered either richness or evenness individually.

### The association between microbiome dominance and disease strengthens in regionally-matched data

The correlation we saw between microbiome dominance and disease persisted in a regionally-matched comparison between disease and microbiome data, and therefore is unlikely to be driven by biogeographic confounders. While the trend between microbiome dominance and coral disease is compelling across our full dataset, not all coral diseases are cosmopolitan and some exist in only one or a few locations (33). As mismatches between region and disease biogeography could confound our overall results, we sought to assess whether large-scale regional effects drive this trend. For example, perhaps high-dominance corals happen to live in high-disease areas, resulting in incidental correlations between dominance and disease. To test for regional effects, we repeated the PGLS analyses restricting the data to only coral microbiomes from Australia, where sampling was most intensive and for which we have long-term disease datasets best-matched to the microbiome data. In this analysis, ecological dominance in Australian coral tissue microbiomes predicted disease prevalence even more strongly under the lowest AICc model (PGLS: R^2^ = 0.49, p = 0.00015, FDR q = 0.005). However, this correlation was strong under all models (Supplementary Data Table 3b). A likely explanation for this stronger result is simply that the disease and microbiome data were drawn from the same region in this analysis, whereas in other cases the available disease and microbiome data were only partially regionally matched. These stronger results in the Australia-only model suggest that microbiomes vary enough geographically that disease and microbiome data from the same location produce the clearest correlations.

### Beta diversity explains little variation in disease susceptibility

Animal microbiomes are often conceived of as having some compositions that are associated with health, and others that are dysbiotic or unhealthy. We sought to test whether this same microbiome beta-diversity framework could predict the extent to which healthy members of different coral taxa are vulnerable to disease. To do so, we correlated coral disease susceptibility against the top three principal coordinate (PC) axes from Weighted and Unweighted UniFrac analyses of microbiome beta-diversity. In contrast to the strong association between microbiome dominance and disease, microbial community composition had less pronounced associations with disease susceptibility. Weighted UniFrac PC axis 3 only nominally significantly correlated with disease susceptibility in all compartments, but this relationship did not remain significant after accounting for multiple comparisons (PGLS: R^2^ = 0.26, p = 0.04, FDR q = 0.90; Supplementary Data Table 4).

### Microbiome dominance vs. disease correlations are driven by γ-proteobacteria

Ecological dominance itself seems an unlikely structural property to act as a mechanism of disease resistance. Therefore, we investigated if this high-level summary measure reflected the effects of some specific microbe or set of microbes. For example, disease susceptibility among *Acropora* has been shown to correlate with the abundance of *Rickettsiales* in coral tissues (34, 35).

To test how shifts in the dominant class of microbes in coral tissue interacted with the dominance-disease correlation, we repeated our previous correlations twice: once in coral genera that are α-proteobacteria dominated, and once in coral genera that are γ-proteobacteria dominated. Both datasets were visualized with ancestral state reconstruction (Supplementary Figures 2a and 2b). Correlations between microbiome dominance and disease were visually apparent only in reconstructions of the γ-proteobacteria dominated corals, and the dominance-disease correlation was far stronger in γ-proteobacteria dominated corals (PGLS: R^2^ = 0.50, p = 0.0001, FDR q =0.003; Supplementary Data Table 3c), where dominance explained most (50%) of the variation in disease susceptibility. In contrast, α-proteobacteria dominated tissue microbiomes showed no discernable dominance-disease correlations either visually or statistically (PGLS: R^2^ = 0.06, p = 0.31, FDR q = 0.81; Supplementary Data Table 3c). This suggested that overall dominance-disease correlations are unlikely to be driven by α-proteobacteria, but may be driven by γ-proteobacteria or specific taxa within this bacterial class. Critically, nothing about these results contradicts the possibility that some α-proteobacteria are coral pathogens, parasites, or opportunists (36). It merely suggests that in healthy corals, dominance by α-proteobacteria does not predict the overall level of disease susceptibility of coral genera, whereas dominance by one or more γ-proteobacteria does.

### The coral symbiont *Endozoicomonas* drives dominance-disease correlations

Bacteria in the genus *Endozoicomonas* are among the most-studied γ-proteobacterial symbionts of corals. In several species *Endozoicomonas* forms prominent aggregates known as CAMAs (coral associated microbial aggregates) in coral tissue (31). In species where *Endozoicomonas* is common, it frequently decreases in relative abundance during coral bleaching or disease (37), suggesting a commensal or mutualistic rather than opportunistic relationship with host health. Further, it has previously been observed that the family *Endozoicomonadaceae* shows by far the strongest signal of cophylogeny with coral hosts among tested bacterial families in coral tissue (16). In the present dataset, *Endozoicomonas* was also the single genus that most typically dominated coral tissue microbiomes (18% of coral genera; Figure 2A). We therefore tested whether the signal of microbiome dominance on disease susceptibility could be explained by the abundances of dominant taxa, and found that across all corals in our dataset, *Endozoicomonas* abundance explained the overwhelming majority of variation in ecological dominance among coral tissue microbiomes (PGLS: R^2^: 0.60, p = 6.2 ×10^−10^, FDR q = 2.5×10^−9^; Figures 2C & 3A; Supplementary Data Table 5a). Further, the relative abundance of *Endozoicomonas* in coral tissue alone explained 30% of variance in overall disease susceptibility (PGLS: R^2^ = 0.30, p = 0.0002, FDR q = 0.0004; Figure 3B; Supplementary Data Table 5a), exceeding the signal from ecological dominance.

**Figure 3.**
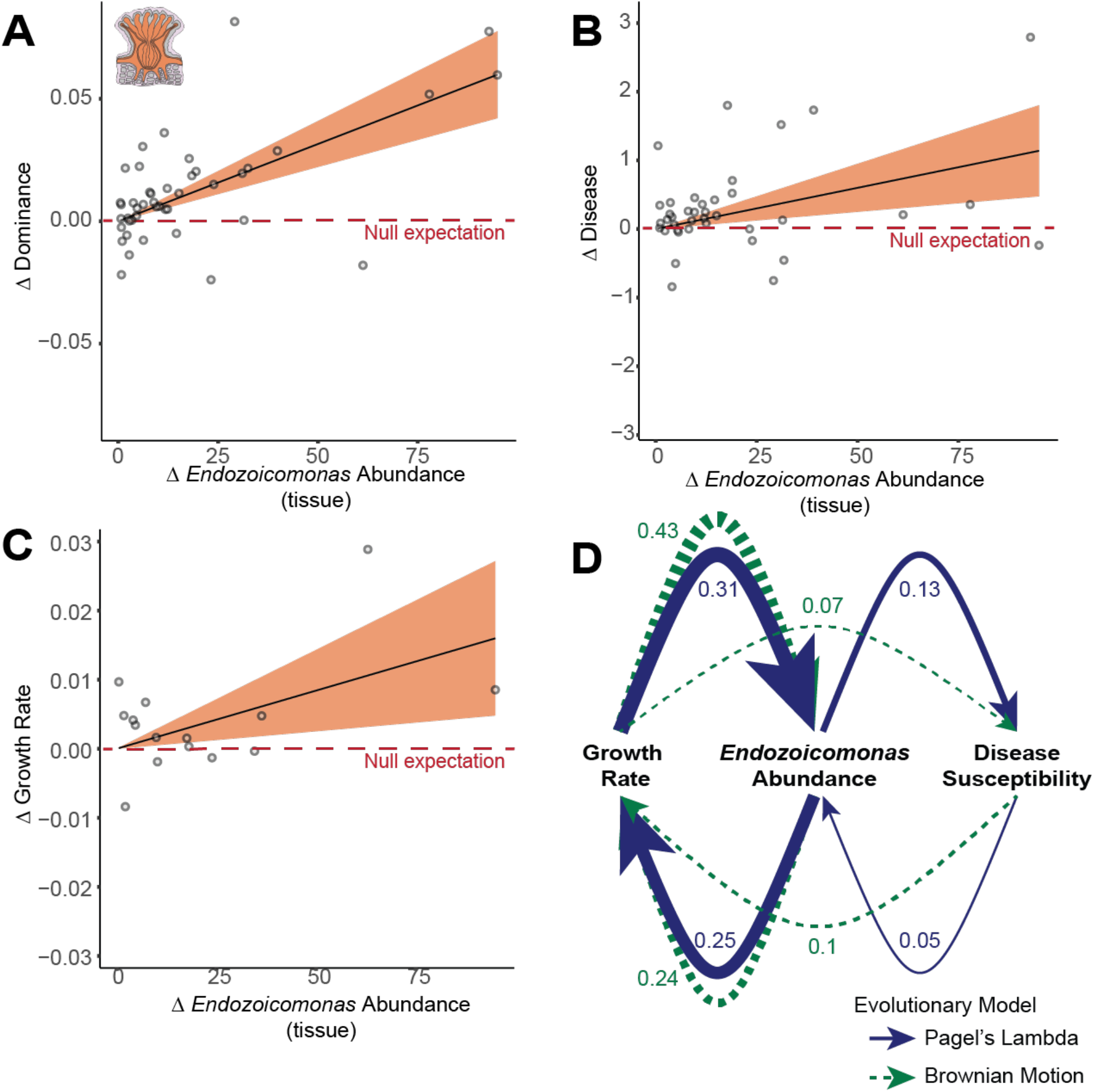
*Endozoicomonas* correlates with growth and disease. Phylogenetic independent contrasts in *Endozoicomonas* relative abundance in coral tissue (per 1000 reads) against **A)** microbial dominance, **B)** integrated estimate of coral disease prevalence and **C)** coral growth rate (mm per year) from the coral traits database. Dotted red lines in panels B and C indicate the null expectation that if traits are uncorrelated, change in the x-axis trait will not correlate with changes in the y-axis trait, with contrasts distributed equally above or below the dotted line. Statistics reflect phylogenetic generalized least squares (PGLS) regression (Supplemental Data Tables 5 and 9). **D)** Modeled direction of causality between *Endozoicomonas* abundance, disease susceptibility and growth rate using both Brownian Motion (blue) and Pagel’s Lambda (green, dotted) evolutionary models. The thickness of the lines represents the averaged standardized path coefficients of the top competing models based on CICc values (Supplementary Data Table 11).

After testing *Endozoicomonas -* disease associations as a prior hypothesis, we also sought to put these associations in context by testing for correlations with disease in all other dominant microbial genera found in the study (Figure 2C; Supplementary Data Table 6a). This scan confirmed that *Endozoicomonas* showed far stronger correlations with disease than other microbes. It also found two dominant genera found in coral skeleton that also significantly correlated with disease susceptibility: *Cellulosimicrobium* (Phylum: Actinobacteria/Actinomycetota) and *Paramaledivibacter* (Phylum: Firmicutes/Bacillota), but to a much lesser extent than *Endozoicomonas* in tissue and mucus compartments. Thus, prior results linking ecological dominance and overall disease susceptibility appear to be largely explained by changes in *Endozoicomonas* relative abundance over coral evolution.

### Coral opportunist abundance in healthy corals does not predict genus-wide disease susceptibility

Correlations between *Endozoicomonas* and disease across the coral tree were initially surprising, as *Endozoicomonas* is not thought to be associated with coral pathogenesis. This raised the question of whether the abundance of known or suspected coral pathogens in apparently healthy corals correlates with cross-genus differences in disease susceptibility.

The abundance of bacterial groups containing prominent putative bacterial pathogens (such as *Vibrionales, Nostocales* or *Rickettsiales*, see (38)) in healthy corals did not show any correlation with disease susceptibility among coral species when tested (Supplementary Data Table 7). Thus, having high abundances of coral opportunists when healthy does not seem to be a hallmark of disease-susceptible corals. This is mostly expected since the abundance of pathogens typically only increases during stress. These observations in healthy corals leave open the question of what about *Endozoicomonas* causes it to be so strongly correlated with coral disease susceptibility.

*Endozoicomonas* is linked to metabolic benefits to the coral host (39, 40) and experimental studies have shown that decreases in its abundance is typical with disease (41, 42) or other health stressors such as bleaching (37). This suggests that the striking correlation between *Endozoicomonas* and disease is not due to pathogenesis by *Endozoicomonas*, but instead might arise due to opportunity costs (e.g., in innate immunity, permissiveness to CAMA formation, or symbiosis with defensive microbes within coral tissue). If maintenance of high abundances of *Endozoicomonas* has fitness costs, they may be balanced by metabolic benefits, and we should expect that *Endozoicomonas* would be more abundant in corals with life-history strategies that favor traits such as rapid growth.

### *Endozoicomonas* is associated with high growth rates

If symbiosis with *Endozoicomonas* did play a causal role in coral life-history tradeoffs, we hypothesized that we would see a positive correlation between a beneficial coral trait and *Endozoicomonas* that counterbalances the correlation between *Endozoicomonas* and disease. Given that *Endozoicomonas* is thought to be a metabolic mutualist of corals, and it has recently been suggested to facilitate faster coral growth (43), growth rate seemed like a likely candidate for a potential benefit explaining the persistence of coral-*Endozoicomonas* associations. Depending on the mechanism of action, any such *Endozoicomonas* - growth correlations might depend merely on the presence of *Endozoicomonas*, or alternatively on its relative abundance. Using data from the Coral Trait Database (CTDB; (28)) we tested whether *Endozoicomonas* relative abundance was correlated with growth rate in corals where we detected *Endozoicomonas* (i.e., the effect of relative abundance alone) and in all corals (i.e., the combined effect of presence and abundance). In both cases, we limited this analysis to only corals with replicated growth rate data (>= 5 replicates in the CTDB).

While the abundance of *Endozoicomonas* was not correlated with growth rate across all coral genera (tissue PGLS: R^2^ = 0.11, p = 0.17, FDR q = 0.17; Supplementary Data Table 8a), across coral genera where *Endozoicomonas* was detected (n = 17 genera), its relative abundance was strongly correlated with growth rate (tissue PGLS: R^2^ = 0.31, p = 0.024, FDR q = 0.024; Supplementary Data Table 8b). These results are consistent with a pattern in which lineage-specific expansions of *Endozoicomonas* within coral microbiomes correlate with or potentially contribute to growth rate. Thus, *Endozoicomonas* may in part explain, or at least correlate with, about a third of known growth rate differences between coral genera.

Thus, across the coral genera surveyed in our dataset, initial, low-level symbiosis with *Endozoicomonas* does not correlate with growth rate, but subsequent expansions of the abundance of *Endozoicomonas* within coral microbiomes co-occur with both higher average growth rates and greater disease susceptibility.

### *Endozoicomonas* may mediate growth-defense tradeoffs during coral evolution

Having seen that *Endozoicomonas* is correlated with both disease susceptibility and growth-rate in corals, we investigated if these correlations were stronger or weaker than the direct correlation between disease and growth rate in our dataset. Across genera with both growth rate and disease prevalence data, growth and disease susceptibility were positively correlated. However, this correlation had only a modest effect size and was not statistically significant. Thus, in this dataset *Endozoicomonas* showed stronger associations with both growth and disease than these factors showed with one another, regardless of whether the analysis was conducted across all coral genera (tissue PGLS: R^2^ = 0.12, p = 0.17, FDR q = 0.17; Supplementary Data Table 9a) or just those where *Endozoicomonas* was present (tissue PGLS: R^2^ = 0.06, p = 0.37, FDR q = 0.37; Supplementary Data Table 9b). This suggested that *Endozoicomonas* relative abundance might not merely mark tradeoffs between growth and disease, but may play some causal role in one or both processes.

### Phylogenetic path analysis of growth, disease, and *Endozoicomonas* abundance

The univariate correlations between *Endozoicomonas*, host disease susceptibility and growth rate raise the question of the direction of causality by which these factors have become non-randomly associated during coral evolution. Using phylogenetic path analysis (Methods), we compared 14 models of the relationship between *Endozoicomonas* relative abundance, disease susceptibility, and growth rate (Supplementary Data Table 10a, Supplementary Figure 3).

As is common in this type of analysis, more than one model was consistent with the data. However, none of the top models using either BM (Supplementary Table 10b) or Pagel’s lambda (Supplementary Data Table 10c) suggested that disease influenced growth rate or vice versa without the influence of *Endozoicomonas* (Figure 3D), and all significant models include *Endozoicomonas*. Thus, while the precise feedback remains to be determined, causality analysis suggests that, in some capacity, *Endozoicomonas* likely mediates growth rate and disease.

### Potential mechanisms of action

The findings of positive correlations between *Endozoicomonas*, host growth rate, and host disease susceptibility documented in this study complement and contextualize much of the ongoing work on the mechanisms underlying proposed coral-*Endozoicomonas* metabolic mutualism (39, 43) and suggest that the interaction of *Endozoicomonas* with coral disease susceptibility deserves greater scrutiny. They also echo findings of correlations between life-history strategy and microbiome structure in other important marine invertebrates, such as that between predator defense and microbial abundance in marine sponges (44).

The mechanism by which corals with high proportions of *Endozoicomonas* become more vulnerable to disease are not yet known, but potential explanations fall into three main categories: ecological, structural, or immunological.

Many coral microbes (but not *Endozoicomonas*) are thought to protect against pathogenic disease by mechanisms such as antibiotic secretion (24), direct predation (25), jamming of quorum signaling (26), and through physically occupying space close to host tissues that may restrict binding sites for opportunists and pathogens. In theory, it is possible that *Endozoicomonas* abundance may interact with other aspects of coral microbial ecology, thereby reducing microbially-derived host defenses. However, that *Endozoicomonas* are frequently observed in discrete CAMAs complicates this possibility, as any effects on microbes outside the local area of these CAMAs would have to rely on indirect consequences of *Endozoicomonas*-coral interactions or secreted factors. Nevertheless, if this hypothesis were correct, the reductions in the abundance of *Endozoicomonas* that are often reported in diseased coral phenotypes (e.g., (37)) would then be adaptive on the part of the host, by allowing proportionally greater growth of other, more protective microbes. This hypothesis could be tested by microbial inoculation experiments that increase *Endozoicomonas* abundances prior to or concurrent with disease exposure, with the prediction that this would increase disease severity (although care must be taken to exclude nutritional benefits from corals directly eating the *Endozoicomonas* confounding the results). More systematic studies of whether high abundances of *Endozoicomonas* are exclusively found in visible CAMAs could also speak to the plausibility of this ecological hypothesis, by clarifying the likely routes for interaction between *Endozoicomonas* and other coral-associated microbes.

In addition to ecological interactions, the *Endozoicomonas* - disease susceptibility correlation may also arise as a result of host traits that are permissive for the formation of microbial aggregates. As the cellular processes involved in establishing mutualism, commensalism and pathogenesis often overlap, the same host-microbe interactions that allow *Endozoicomonas* and some other microbes like *Simkania* (43) to aggregate within coral tissues may also be more permissive towards invasion by pathogens. So far known coral pathogens have not been reported to be present within CAMAs. However, other structural mechanisms are possible. For example, the density, morphology, or diversity of septate junctions — which form epithelial barriers similar to tight junctions in chordates (45) — might, in theory, influence the ability of both *Endozoicomonas* and pathogenic microbes to enter coral tissues. This idea could be tested by examining cellular morphology, sequence similarity, and/or gene expression of septate junctions and their constituent components in coral species in which CAMAs did or did not form.

Finally, it is possible that coral immunological strategies that permit symbiosis with high abundances of *Endozoicomonas* also tend to make corals more vulnerable to pathogens. Coral species vary in immune investment (as measured by immune parameters like melanin abundance, phenoloxidase activity, etc), and low immune investment has been observed to correlate with disease susceptibility (46). Some theory predicts that the evolution of more permissive immunological strategies is favored by symbionts that provide metabolic benefits to the host (47). In corals specifically, immune repertoires in key gene families such as TIR-domain containing genes vary greatly between species, which has been hypothesized to influence microbiome structure (48). Thus, symbiosis with *Endozoicomonas* may promote lower immune investment in corals, which in turn increases disease susceptibility. This hypothesis could be tested by comparing the length of coral-*Endozoicomonas* associations, to see whether longer histories of association lead to low immune investment, or by examining selection on innate immune genes in low vs. high *Endozoicomonas* coral lineages (e.g., by dN/dS ratios).

A related immunological explanation would occur if *Endozoicomonas* itself achieves high abundances by suppressing aspects of host immunity. Genomic studies of host-associated *Endozoicomonas* identified variation in the proportion of eukaryote-derived genes and domains as a key feature of strain variation, including some domains thought to suppress immunity-induced apoptosis (49). If representatives of those different strains could be cultured, experiments adding exogenous *Endozoicomonas* might clarify whether *Endozoicomonas* strains have any direct effects on coral immunity, and if so whether they differ from strain to strain.

## Conclusions

Animals evolved in a microbial world resulting in interactions between animal hosts and their associated microbes that influence organismal fitness. These interactions can occur across generations and may elucidate many of the eco-evolutionary patterns that we see among organisms, including mediation of important life history tradeoffs. Using evolutionary analyses of coral microbiomes, we provide evidence that symbiosis with *Endozoicomonas* may mediate growth vs. disease resistance (defensive) tradeoffs. While further manipulative studies are necessary to confirm this finding and determine the directionality of the relationship, evidence for this trend across the coral tree of life is compelling.

Our comparative approach suggests that *Endozoicomonas*-dominated lineages of corals may grow more quickly under ideal conditions but are more likely to succumb to coral disease. Because much other work has shown that coral disease is exacerbated by global and local stressors such as climate-change driven heat waves or local pollution events (33, 38), this may make *Endozoicomonas-* dominated coral especially vulnerable to environmental change (Figure 4).

**Figure 4.**
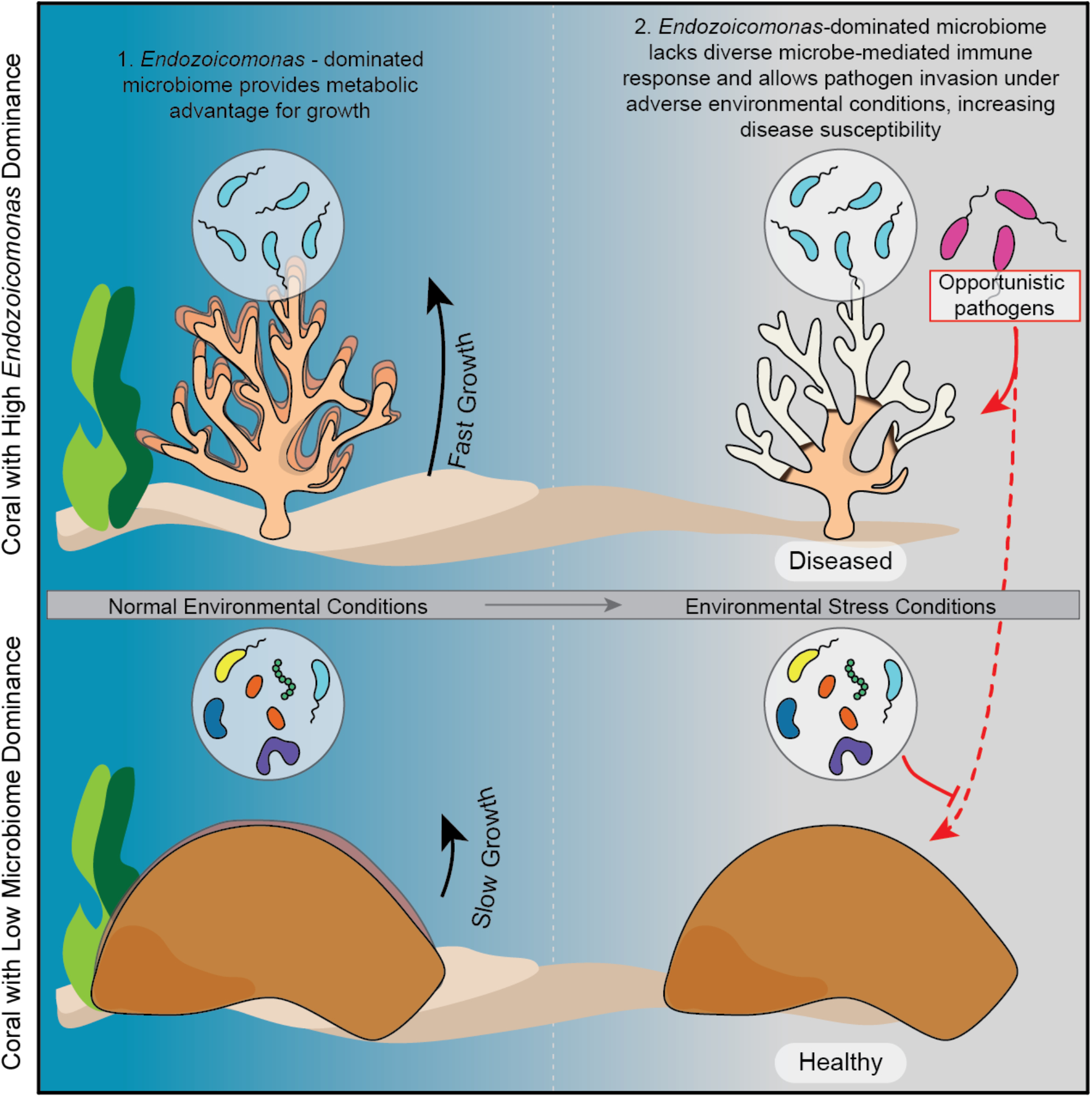
Endozoicomonas dominance facilitates life history tradeoffs. Conceptual hypothesis on the role *Endozoicomonas* (in teal) plays in the tradeoff between growth and defense (disease susceptibility).

If microbial symbiosis does play a causal role in coral life history tradeoffs in the present day, then identifying microbes underlying those tradeoffs may benefit microbiome manipulation for targeted coral conservation and restoration strategies. While the correlation between *Endozoicomonas* and disease in this work was observed at the genus level (primarily because this is the level of taxonomic specificity for most available disease surveys), future work could examine whether similar trends appear between coral sister species or within coral populations. For example, microbial screening (e.g., (50)) could help identify *Endozoicomonas*-dominated coral species or populations that may be more susceptible to disease and drive the conservation and protection of these individuals or their habitats. Identifying these target corals is perhaps most relevant for coral restoration initiatives that include breeding, nursery propagation and outplanting, where coral health is monitored closely and predicting disease susceptibility can inform decision-making. Depending on the mechanism underlying the *Endozoicomonas-*disease susceptibility correlations reported here, *Endozoicomonas*-dominated corals may further represent strong candidates for microbiome engineering (e.g., human-assisted manipulation of host-associated microbes (51) or the application of probiotics (14, 52)) to enhance host resilience in anticipation of stress events by decreasing microbiome dominance. That said, we emphasize that microbiome manipulation and other restoration initiatives are not replacements for efforts to decarbonize global economies to limit greenhouse gas emissions.

The results presented here provide the first evidence of a likely microbe-mediated life-history tradeoff in scleractinian corals. Further exploration of this and other such potential tradeoffs may shed light on the evolutionary interplay between microbes and the physiology and ecology of their animal hosts.

## Methods

### Coral sample collection and 16S rRNA pre-processing

16S rRNA sequence data was obtained from visibly healthy coral DNA extractions collected and processed for the Global Coral Microbiome Project (GCMP). This included coral samples taken from eastern and western Australia that were used in a previous study by Pollock and co-authors (16) in addition to coral samples taken from the Red Sea, Indian Ocean, Coral Triangle, Caribbean, and Eastern Pacific. All samples compared in this study were collected, processed, and sequenced using consistent protocols as outlined below. In total, 1,440 coral, outgroup, and environmental samples were collected. Of these GCMP samples, the 1,283 scleractinian coral and outgroup samples were used in the present study (Supplementary Data Table 1a). These comprise 132 species and 64 genera of corals originating from 42 reefs spanning the Pacific, Indian, and Atlantic oceans.

The collection and processing of these coral samples followed the methods outlined in Pollock et al (16) and are compatible with samples processed for the Earth Microbiome Project (53). Briefly, three coral compartments were targeted for each sample: tissue, mucus, and skeleton. Mucus was released through agitation of coral surface using a blunt 10mL syringe for approximately 30 seconds and collected via suction into a cryogenic vial. Small coral fragments were collected by hammer and chisel or bone shears for both tissue and skeleton samples into sterile WhirlPaks (Nasco Sampling, Madison, WI). All samples were frozen in liquid nitrogen on immediate return to the surface prior to processing. In the laboratory, snap frozen coral fragments were washed with sterile seawater and the tissue was separated from skeleton using sterilized pressurized air at between 800-2000 PSI. Tissue and skeleton samples were then preserved in PowerSoil DNA Isolation kit (MoBio Laboratories, Carlsbad, CA; now Qiagen, Venlo, Netherlands) bead tubes, which contain a guanidinium preservative, and stored at -80° to await further processing. Outgroup non-scleractinian Anthozoans were also opportunistically collected and stored similarly, including healthy samples of the genera *Millepora* (hydrozoan fire coral), *Palythoa* (zoanthid), *Heliopora* (blue coral), *Tubipora* (organ pipe coral), and *Xenia* and *Lobophytum* (soft corals).

Bacterial and archaeal DNA were extracted using the PowerSoil DNA Isolation Kit (MoBio Laboratories, Carlsbad, CA; now Qiagen, Venlo, Netherlands). To select for the 16S rRNA V4 gene region, polymerase chain reaction (PCR) was performed using the following primers with illumina adapter sequences (underlined) at the 5’ ends: 515F (54) 5′− TCG TCG GCA GCG TCA GAT GTG TAT AAG AGA CAG GTG YCA GCM GCC GCG GTA A −3′ and 806R (55) 5’− GTC TCG TGG GCT CGG AGA TGT GTA TAA GAG ACA GGG ACT ACN VGG GTW TCT AAT −3′). PCR, library preparation, and sequencing on an Illumina HiSeq (2×125bp) was performed by the EMP (53). All raw sequencing data and associated metadata for the samples used in this study are available on Qiita (qiita.ucsd.edu) under project ID 10895, prep ID 3439.

### Sequence assembly, quality control and taxonomic assignment

16S rRNA sequencing data were processed in Qiita (56) using the standard EMP workflow. Briefly, sequences were demultiplexed based on 12bp Golay barcodes using “split_libraries” with default parameters in QIIME1.9.1 (57) and trimmed to 100bp to remove low quality base pairs. Quality control (e.g., denoising, de-replication and chimera filtering) and identification of amplicon sequence variants (ASVs) were performed on forward reads using deblur 1.1.0 (58) with default parameters. The resulting biom and taxonomy tables were obtained from Qiita (CRC32 id: 8817b8b8 and CRC32 id: ac925c85) and processed using a customized QIIME2 v. 2020.8.0 (59) pipeline in python (github.com/zaneveld/GCMP_global_disease). Taxonomic assignment of ASVs was performed using vsearch (60) with SILVA v. 138 (61).

### Removal of cryptic mitochondrial reads

Coral mitochondrial reads obtained from metaxa2 (62) were added to the SILVA repository to better identify host mitochondrial reads that may be present in the sequencing data (63). We refer to this expanded taxonomy as “silva_metaxa2” in code. After taxonomic assignment, all mitochondrial and chloroplast reads were removed. The bacterial phylogenetic tree was built using the SATé-enabled phylogenetic placement (SEPP) insertion technique with the q2-fragment-insertion plugin (64) to account for the short-read sequencing data, again using the SILVA v. 138 (61) database as reference taxonomy. The final output from this pipeline consisted of a taxonomy table, ASV feature table and phylogenetic tree that were used for downstream analyses.

### Identification of potential contaminants

Potential contaminants from extraction and sequence blanks (n = 103 negative controls) were identified and removed using the decontam package (65) in R v. 4.0.2 (66) with a conservative threshold value of 0.5 to ensure all ASVs that were more prevalent in negative controls than samples were removed (n = 662 potential contaminants). The final feature table consisted of a total of 1,383 samples, 195,684 ASVs, and 37,469,008 reads.

### Summary of disease data by coral genus

Disease data were gathered from long-term multi-species surveys in the Florida Keys (the Florida Reef Resilience Program (FRRP), https://frrp.org/), Hawai’i (HICORDIS (29)), and Australia (this study). Disease counts for Australian corals were collected over a period of 5 years (2009-2013) across 109 reef sites and 65 coral genera (Supplementary Data Table 1b). At each of the 109 reefs, we surveyed coral health using 3 replicate belt transects laid along reef contours at 3-4m depth and approximately 20m apart using globally standardized protocols (67). Depending on the reef location, belt transects were either 10, 15, or 20m in length by 2m width making the area surveyed at each reef between 60 and 120m^2^. Within each belt transect, we identified each coral colony over 5 cm in diameter to genus and classified it as either healthy (no observable disease lesions) or affected by one or more of six common Indo-Pacific coral diseases (according to (68)). Together with the FRRP and HICORDIS data, the combined disease dataset contained 582,342 coral observations across 99 coral genera (Supplementary Data Table 1c).

Because many of these disease observations identified corals only to genus, disease prevalence data were summarized at the genus level. All three resources represent coral surveys over time, ranging from 5 to 16 years. We chose such long-term datasets in an attempt to minimize the potential effects of specific events (e.g., bleaching in a single summer) and instead to capture more general trends in disease susceptibility across species, if such trends were present. Summarizing these data at the genus level was thus part of a comparative strategy, enabling us to extract overall trends and average out local circumstances, so that we could find holobiont features that control disease resistance that may protect some corals but not others.

When summarizing at the genus level, individual counts of healthy corals or corals with specific diseases were summed within coral genera across these datasets.

To ensure sufficient replication, we excluded coral genera with fewer than 100 observed individuals. This minimal count was selected because it is the lowest frequency at which diseases with a reasonably high frequency (e.g., 5%) can be reliably detected. (With 100 counts, there is a >95% chance of detecting at least one count of any disease present with >= 5% prevalence; cumulative binomial, 100 trials, success chance = 0.05). Because only very rarely observed taxa were removed, this filtering preserved 99.8% of total observations. Ultimately, our genus-level summary produced a table with 581,311 observations across 60 coral genera (Supplementary Data Table 1d).

### Summary of the microbiome data by coral host genus

Statistical summaries of microbiome community composition were calculated for each sample in QIIME2 (59), and then summarized within anatomical compartments and coral genera. These summaries of coral microbiome alpha diversity were richness (observed features per 1000 reads), evenness (the Gini Index), and Simpson’s Index, which combines both richness and evenness. Thus, each combination of coral genus and anatomical compartment — such as *Acropora* mucus — was assigned an average α-diversity value.

Simpson’s Index, which is of particular importance in these results, is at its highest when a single taxon is the only one present in microbiome, and at its lowest when there are both a large number of taxa, and all taxa have equal abundance. Thus, this measure is reduced both by community richness and community evenness (Simpson’s Index is closely related to Simpson’s Diversity, which is calculated as 1 - Simpson’s Index, such that more rich or even communities produce higher values).

### Construction of a genus level trait table

The summarized, genus-level disease susceptibility data compiled from all disease projects, and the summarized genus-level microbiome diversity data (see above) were combined to form a trait table that was used in subsequent evolutionary modeling. Additionally, the relative abundance of ‘dominant’ microbes analyzed in this study was averaged within genera and added to this genus-level trait table.

### Genus-level summary of a reference coral phylogeny

Starting with a previously published multigene time-calibrated phylogeny of corals (30) that we had previously used to demonstrate phylosymbiosis in corals (16), we randomly selected one representative species per genus to produce a genus level tree. This approach was preferred over several alternatives — such as trimming the tree back to the last common ancestor of each genus and reconstructing trait values — because it required fewer assumptions about the process of trait evolution. As microbiome data were not available for all genera on the coral tree (e.g., temperate deep sea corals), the tree was further pruned to include only the subset of branches that matched those with microbiome data.

### Addition of genus-level coral growth data

To examine the influence of microbiome structure on coral traits, we pulled growth data from the Coral Trait Database (28) from all coral genera that matched those with both microbiome and disease data, and were collected using consistent metrics (mm/yr). This resulted in growth rate data from 18 coral genera that were subsequently combined with our genus-level trait table (Supplementary Data Table 1d).

### Phylogenetic Correlative Analysis

Shared evolutionary history induces correlations in traits between species that violate the requirement of standard statistical tests that observations must be independent and uncorrelated. Thus, special care must be taken to account for phylogeny in comparative analysis. We first applied Felsenstein’s phylogenetic independent contrasts (PIC) to visualize our cross-genus trait correlations using the phytools R package (69). This method removes the effect of any shared evolutionary histories by calculating differences in trait values (contrasts) between sister taxa. We next examined the relationships between traits using information-theoretic model selection (that is, comparison of AICc scores) to identify phylogenetic generalized least squares (PGLS) models of evolution that best explained the observed distribution of microbiome α- or β-diversity and disease susceptibility (as continuous evolutionary characters) in extant species. We tested 4 evolutionary models in the caper R package (70). In the first model, we used PGLS with no branch length transformation (i.e. holding λ,***δ***, κ = 1). Thus, this first model is equivalent to PIC. In the next 3 models, we transformed branch lengths on the tree by allowing the model to fit either λ, ***δ***, or κ (see below) using maximum likelihood estimation, while fixing the other 2 parameters at 1. We refer to these 4 models as PGLS, PGLS + λ, PGLS + ***δ***, and PGLS + κ. For detailed explanations of each parameter, please refer to Supplementary Data Table 11.

Typically, these models estimated very low λ (∼0), indicating little or low phylogenetic inertia. R^2^ and *p*-values were adjusted for multiple comparisons using a false discovery rate (FDR) correction. Significant relationships between the two traits suggests that they are evolutionarily correlated. Data were visualized by plotting phylogenetic contrasts and all statistics reported represent the best PGLS model results. Additionally, ancestral state reconstructions of key traits were visualized using the contmap function in the phytools R package (69), which in turn estimates internal states using fast maximum-likelihood (ML) ancestral state reconstruct as implemented in the fastAnc phytools function.

### Phylogenetic causality analysis

Observing that A and B are correlated famously does not guarantee that A causes B. However, non-random correlation between A and B does imply some causal association - though there are many possibilities (A causes B, B causes A, a positive feedback loop exists between A & B, some external factor C causes both A and B, etc.). Path analysis represents hypotheses of causality using directed acyclic graphs, then tests the different strengths of association predicted under different hypotheses of causation to test which are consistent with data. The cross-species nature of these data further necessitated use of phylogenetic path analysis, which also accounts for expected trait correlations among related genera. Hypotheses of the direction of causality between microbiome (specifically *Endozoicomonas*), disease, and growth rate were tested using a phylogenetic causality analysis performed in the R package phylopath (71). This analysis tests the ability of different models to explain correlations in trait data. For example, does selection for a high growth rate in turn drive selection for increased *Endozoicomonas* abundance, which then increases disease susceptibility, or does symbiosis with *Endozoicomonas* itself separately increase disease and growth? Fourteen potential causality models were tested to incorporate all biologically plausible pathways between *Endozoicomonas* abundance, disease susceptibility, and growth rate (Supplementary Data Table 11a; Supplementary Figure 3). The top performing causality models according to CICc values (using both Pagel’s λ and Brownian Motion models of evolution) were averaged for interpretation and visualization.

## Acknowledgements

The authors would like to acknowledge many contributors for their field assistance during collection of Global Coral Microbiome Project data in this manuscript. These include: Tasman Douglass, Margaux Hein, Frazer McGregor, Kathy Morrow, Katia Nicolet, Cathie Page, and Gergely Torda for their field assistance in Australia; Valeria Pizarro, Mateo López-Victoria, Alaina Weinheimer, Claudia Tatiana Galindo Martínez for Assistance in Colombia; Chris Voolstra, Maren Ziegler, Anna Roik, and many others at King Abdullah University of Science and Technology (KAUST) for field assistance in Saudi Arabia; Mark Vermeij, Kristen Marhaver, Pedro Frade, Ben Mueller, and others at CARMABI for field assistance during sampling in Curaçao; Danwei Huang for field assistance in Singapore; Ruth Gates, Katie Barrott, Courtney Couch, and Keoki Stender for field assistance in Hawai’i; and Lyndsy Gazda, Jamie Lee Proffitt, Gabriele Swain, and Alaina Weinheimer for their assistance in the laboratory. The authors also acknowledge the staff of the Coral Bay Research Station, Lizard Island Research Station, Lord Howe Island Marine Park, Lord Howe Island Research Station, the Australian Institute of Marine Science, and RV Cape Ferguson for their logistical support. This work was supported by a National Science Foundation Dimensions of Biodiversity grant (#1442306) to RVT and MM, an in-kind UC San Diego Seed Grant in Microbiome Science grant to JZ, an NSF IoS CAREER grant (#1942647) to JZ, and an NSF Postdoctoral Research Fellowship in Biology (#2006244) to HE.

## Supplementary Data Tables

**Supplementary Data Table 1. Microbiome and disease data. a**. Global Coral Microbiome Project metadata **b**. Disease data for Eastern Australia. **c**. Combined genus-level trait table

**Supplementary Data Table 2. Dominant microbial taxa by coral genus. a**. mucus **b**. tissue **c**. skeleton

**Supplementary Data Table 3. PGLS correlations between microbiome alpha diversity and disease. a**. Alpha diversity vs. host disease for all data **b**. Alpha diversity vs. disease for Australia data only **c**. Alpha diversity vs. host disease in gamma-proteobacteria dominated microbiomes

**Supplementary Data Table 4. PGLS correlations between microbiome beta diversity and disease**.

**Supplementary Data Table 5. PGLS correlations between *Endozoicomonas* relative abundance and** host disease prevalence.

**Supplementary Data Table 6. PGLS correlations between all dominant microbes and a**. host disease prevalence **b**. host growth rate.

**Supplementary Data Table 7. PGLS correlations between known opportunists/pathogens and disease**.

**Supplementary Data Table 8. PGLS correlations between *Endozoicomonas* and host growth rate. a**. Correlation of *Endozoicomonas* relative abundance against host growth rate across all coral genera **b**. Correlation of *Endozoicomonas* relative abundance against host growth rate across all coral genera where *Endozoicomonas* is present.

**Supplementary Data Table 9. PGLS correlations between disease susceptibility and host growth rate. a**. Direct correlation of disease susceptibility and growth rate in all coral genera. **b**. Direct correlation of disease prevalence and growth rate in coral genera where *Endozoicomonas* is present.

**Supplementary Data Table 10. Phylogenetic path analysis. a**. Models tested **b**. Relative weight of each model under Brownian Motion. **c**. Relative weight of each model using Pagel’s Lambda.

**Supplementary Data Table 11. PGLS model explanations**.

## Supplementary Figures

**Supplementary Figure 1. Ancestral state reconstruction of disease susceptibility and alpha diversity metrics. a**. richness, **b**. evenness (Gini index) and **c**. dominance.

**Supplementary Figure 2. Ancestral state reconstruction of disease susceptibility and microbiome dominance**. Filtered to coral hosts dominated by **a**. α-proteobacteria or **b**. λ-proteobacteria.

**Supplementary Figure 3. Causality models tested between *Endozoicomonas* abundance, disease susceptibility and growth rate**.

