## Supplementary figures and images for "Evidence for microbially-mediated tradeoffs between growth and defense throughout coral evolution"

### Supplementary Figure 1a

**Supplementary Figure 1a.** Richness vs. Disease

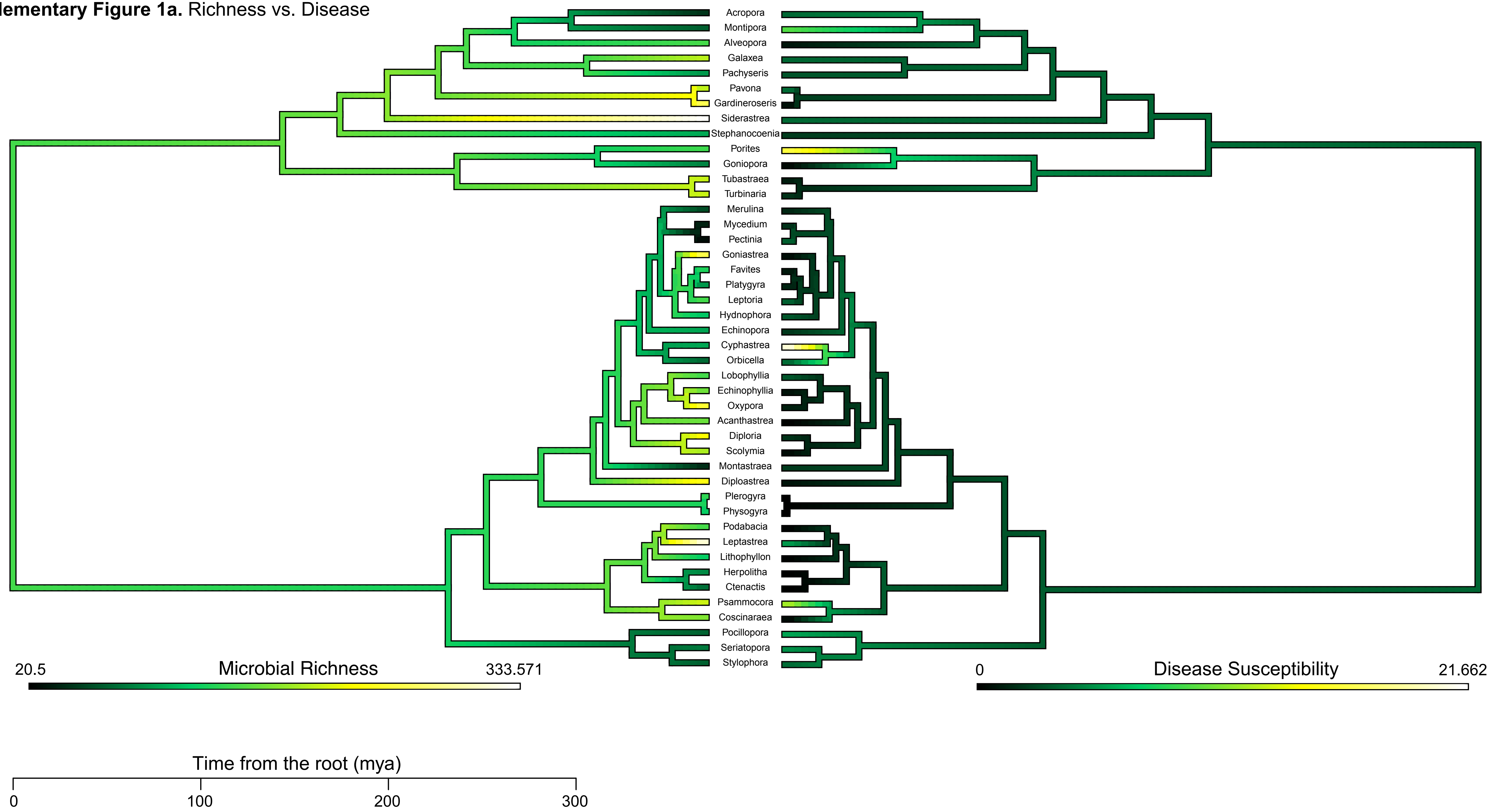

### Supplementary Figure 1b

**Supplementary Figure 1b.** Evenness vs. Disease

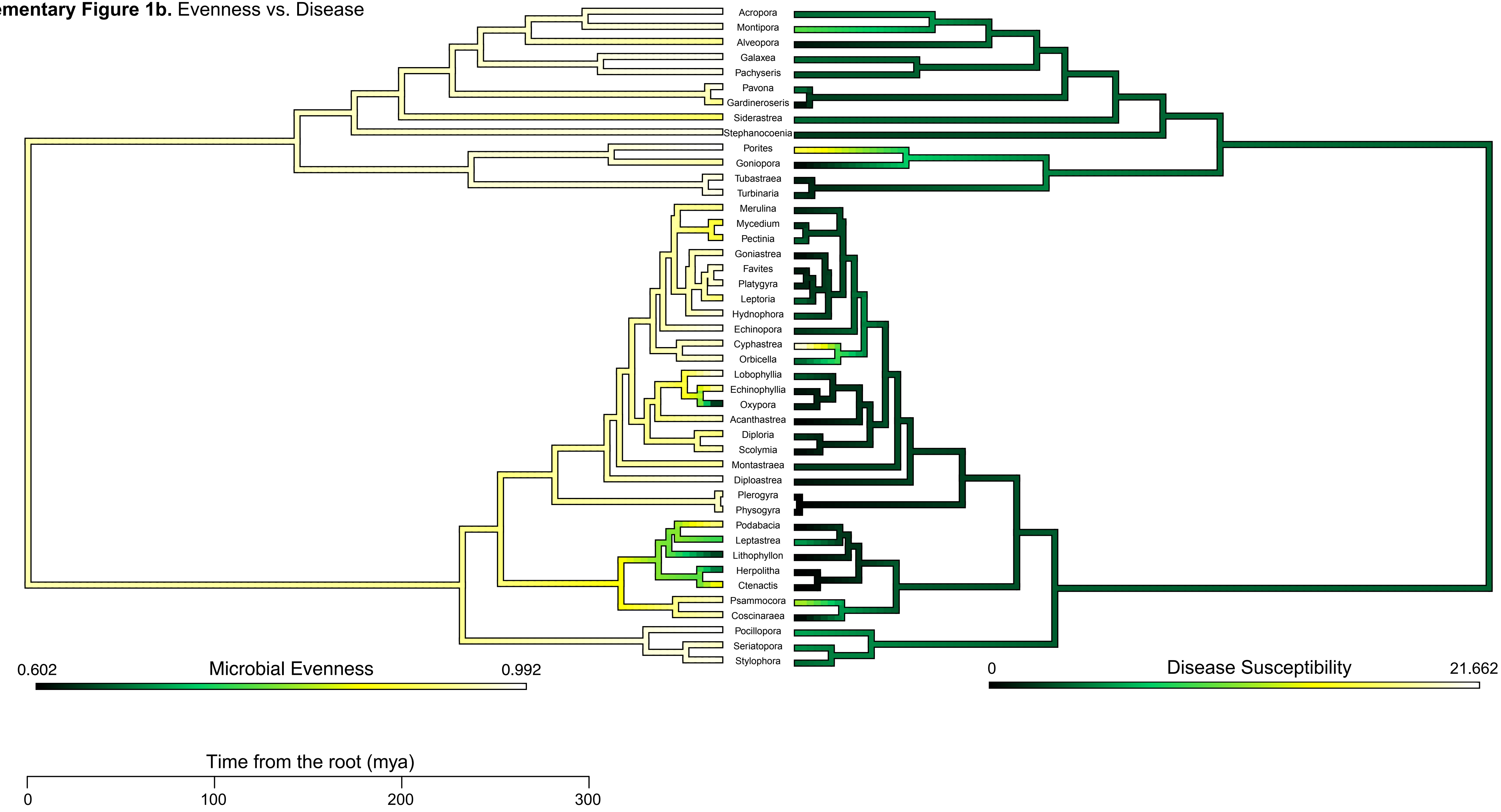

### Supplementary Figure 1c

**Supplementary Figure 1c.** Dominance vs. Disease

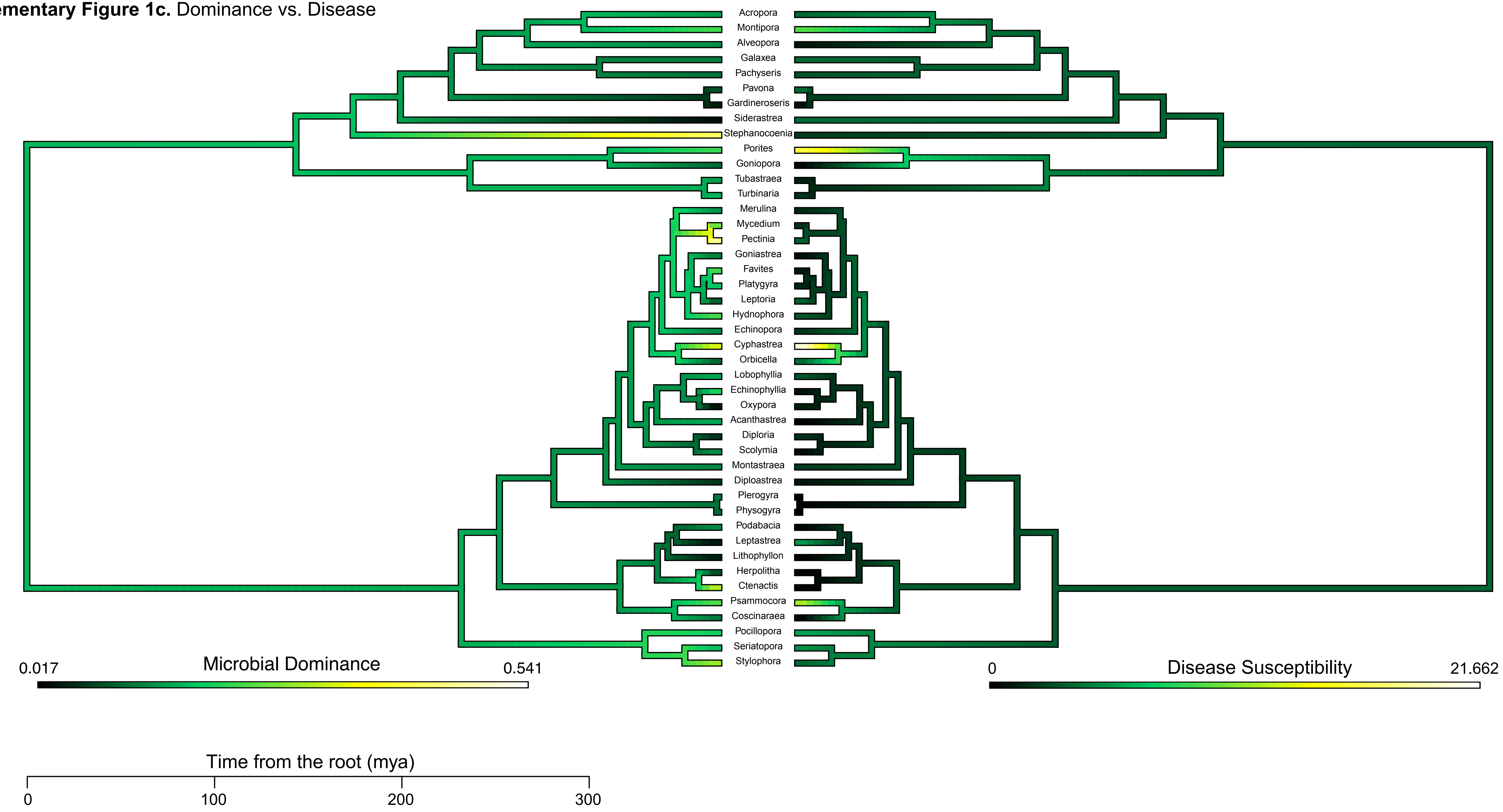

### Supplementary Figure 3

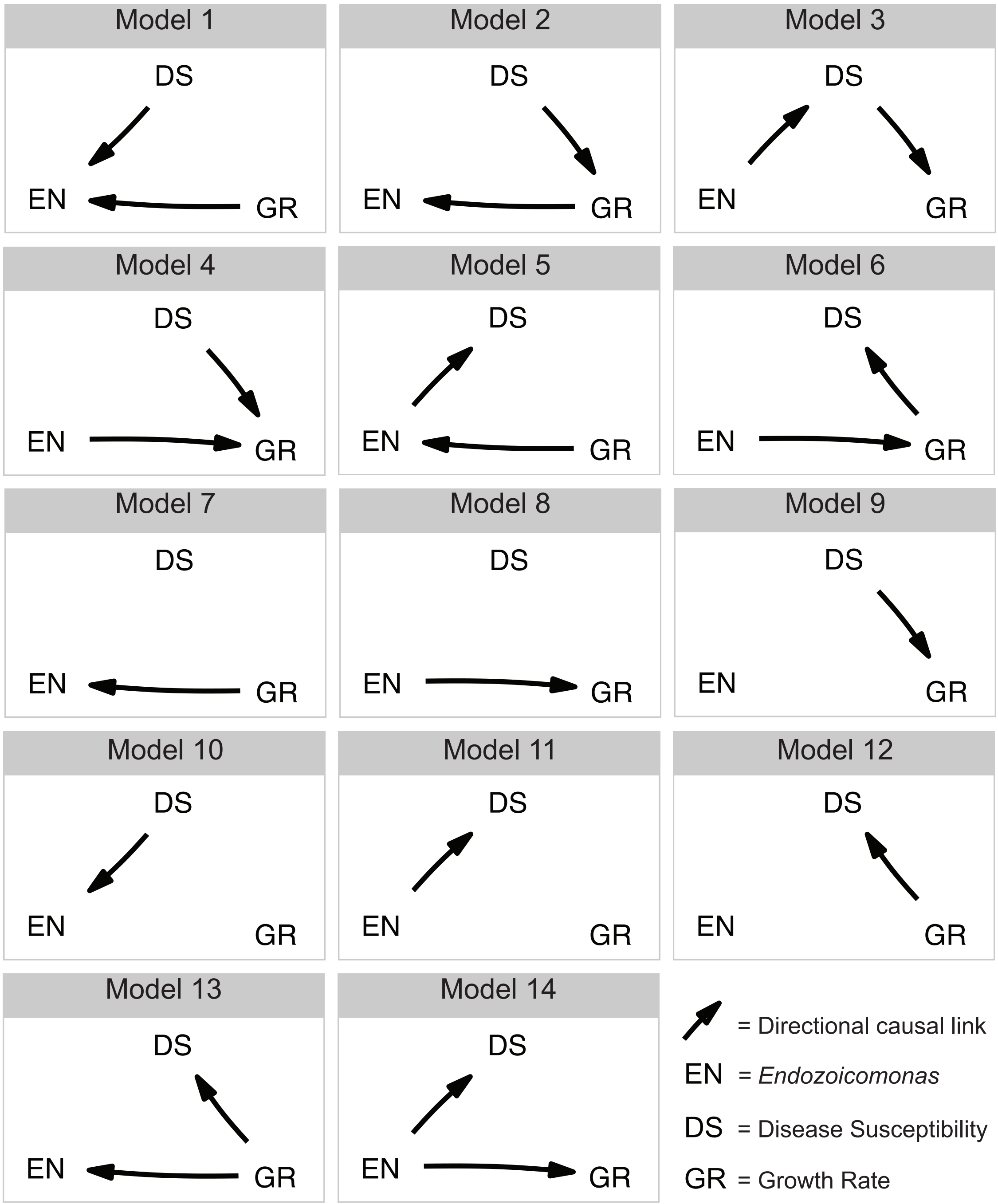
