## Supplementary Figure 2a for "Evidence for microbially-mediated tradeoffs between growth and defense throughout coral evolution"

**Supplementary Figure 2a.** Microbial Dominance (Alphaproteobacteria-dominated hosts) vs. Disease Susceptibility

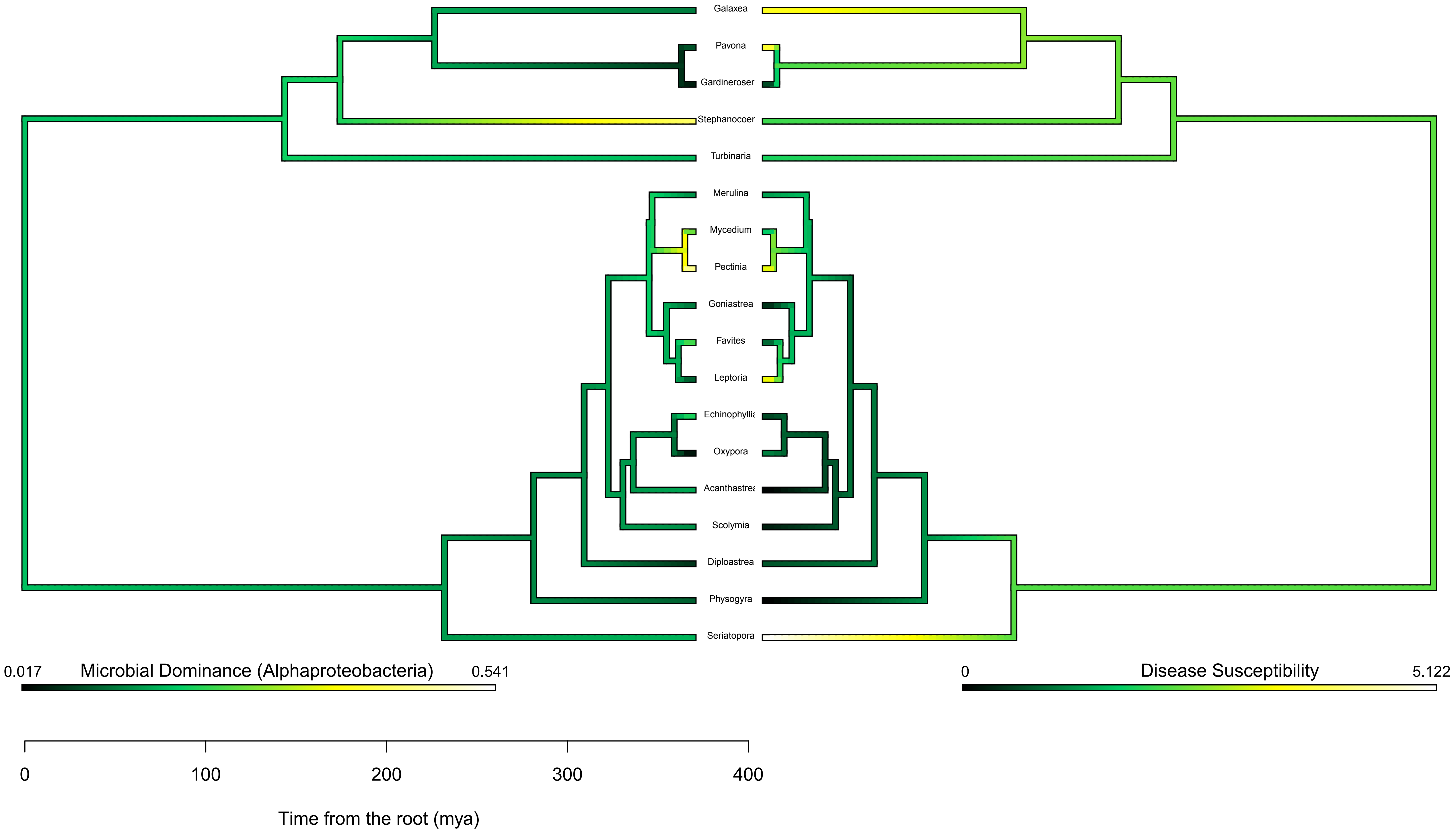
