## Supplementary Figure 2b for "Evidence for microbially-mediated tradeoffs between growth and defense throughout coral evolution"

**Supplementary Figure 2b.** Microbial Dominance (Gammaproteobacteria-dominated hosts) vs. Disease Susceptibility

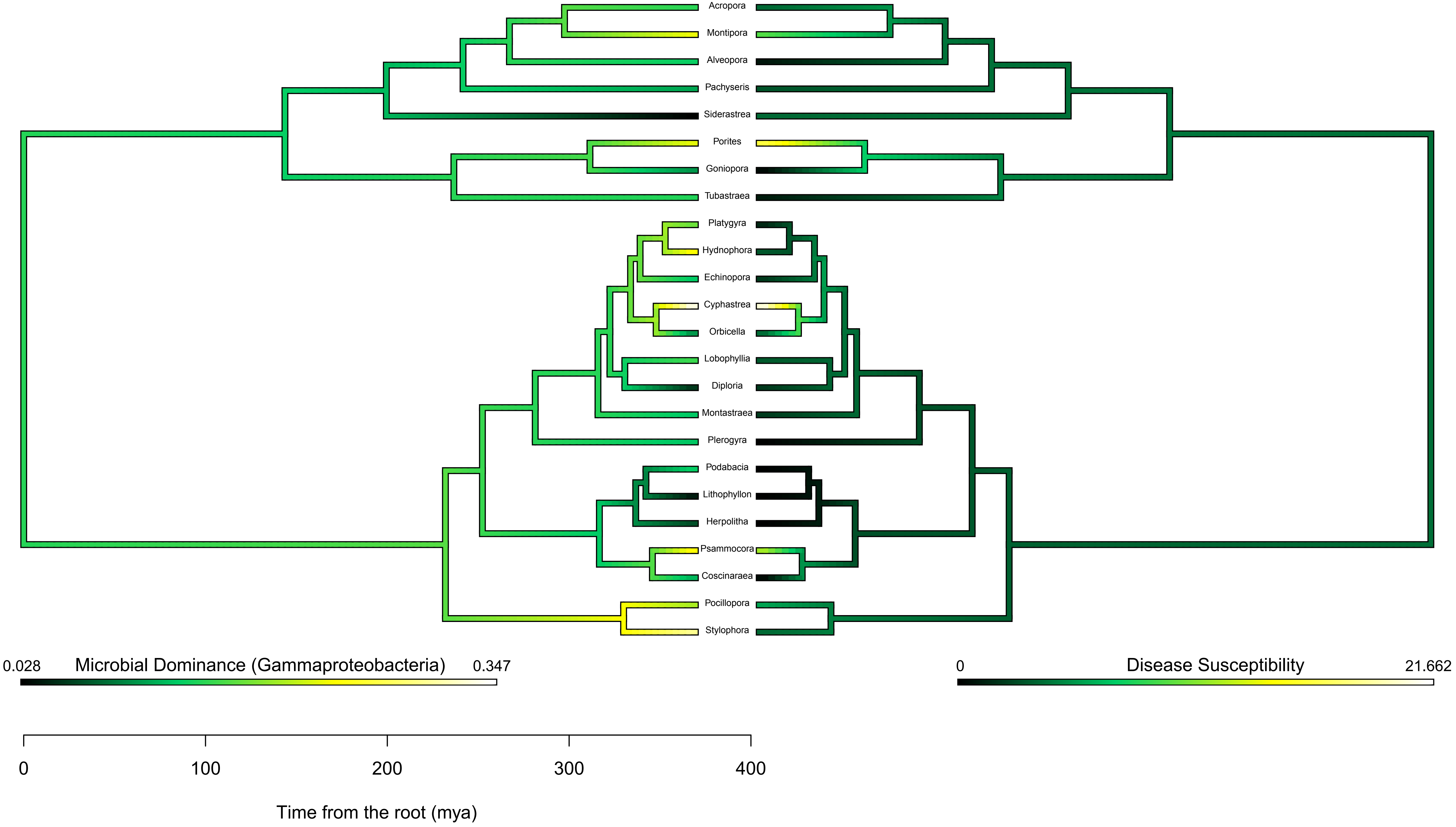
